# Knockdown of PTOV1 and PIN1 Exhibit Common Phenotypic Anti-Cancer Effects in MDA-MB-231 Cells

**DOI:** 10.1101/526319

**Authors:** Shibendra Kumar Lal Karna, Faiz Ahmad, Bilal Ahmad Lone, Yuba Raj Pokharel

## Abstract

**Background:** Earlier, we have identified PTOV1 as a novel interactome of PIN1 in PC-3 cells. This study aims to explore the functional similarity and the common role of both genes in breast cancer cell proliferation.

**Methods:** CTG, crystal violet assay, clonogenic assay, wound healing assay, cell cycle analysis, Hoechst staining and ROS measurement were performed to assess cell viability were performed after knocking down of PTOV1 and PIN1 by siRNAs in MDA-MB-231 and MCF-7 cells. CO-IP, qPCR and western blot were performed for interaction, transcriptional and translational regulation of both genes.

**Results:** Knockdown of PTOV1 and PIN1 inhibited the cell proliferation, colony formation, migration cell cycle, and induces nuclear condensation as well as ROS production. Interaction of PTOV1 and PIN1 was validated by Co-IP in MDA-MB-231 cells. Genes involved in cell proliferation, migration, cell cycle, and apoptosis were regulated by PIN1 and PTOV1. PTOV1 knockdown inhibited Bcl-2, Bcl-xL and induces BAX, LC3 and Beclin-1. Overexpression of PIN1 increased the expression of PTOV1. Knockdown of both genes inhibited the expression of cyclin D1, c-Myc, and β-catenin.

**Conclusions:** PTOV1 and PIN1 interacts and exert oncogenic role in MDA-MB-231 cells by sharing the similar expression profile at transcriptional and translational level which can be a promising hub for therapeutic target.

## 1. Introduction

Breast cancer is the most common cancer occurring in women worldwide (1, 2). Although there are many treatments available like hormone therapy, adjuvant therapy, and surgery, breast cancer remains a major challenge (3, 4). Triple-negative breast cancer (ER, PR, and HER2/Neu negative) cases have poor prognosis and highlight the need to explore the new molecular targets for breast cancer therapy.

Protein-Protein interactions (PPIs) transduce many important cellular functions and their dysregulation can cause diseases. The expression of aberrant proteins seems to enhance their tumor-promoting function due to their interaction with their partners in the cancerous state (5). Identification of cancer enabling PPI hubs that maintain or amplify the cell transformation potential in cancer is one of the major therapeutic strategies in the battle against cancer (6). PIN1 is an established oncogene that regulates the fate of phosphorylated protein catalyzing cis-trans isomerization. PIN1 is overexpressed in breast cancer and mediates its function via RAS signaling, increasing the transcription of c-Jun towards Cyclin D1 (7). Our previous study showed that PIN1 interacts with the novel protein Prostate Tumor Overexpressed 1(PTOV1) in PC-3 cells (8). PTOV1 is a 46 kDa protein with a tandem duplication of two repeated homology blocks of the sequence of 151 and 147 amino acids closely related to each other, located on the 19q 13.3-13.4 chromosome. Chromosome 19 harbors a large number of genes modulated by androgens including PIN1. The Overexpression of PTOV1 in prostate cancer may be due to the cumulative effect of genes residing on chromosome 19. The immunocytochemical analysis of PC-3 cell showed that PTOV1 is located in the cytoplasm close to the nucleus (9). Overexpression of PTOV1 causes the expression of c-Jun both total and in phosphorylated form in prostate cancer cells. PTOV1 interacts with RACK1 to bind with 40s ribosomes during translation initiation stage (10).

The purpose of our study was to reveal how PTOV1 and PIN1 coordinate to drive breast cancer progression, towards this end we used siRNA silencing approach to find out the change in expression profile of various oncogenic signal molecules at transcription and translation levels in MDA-MB-231 cells. Targeting this complex can contribute to autophagy and apoptosis induced cell death increasing the efficacy of the therapeutic approach against breast cancer.

## 2. Methods

### 2.1 Cell line, reagents, and antibodies

MDA-MB-231and MCF-7 breast cancer cell lines were purchased from NCCS, Pune, India. Lipofectamine RNAiMAX and Opti-MEM media were obtained from Invitrogen Corp (Carlsbad, CA, USA). siRNAs were purchased from Qiagen (Hilden, Germany). SYBR Green was obtained from Bio-Rad (Hercules, California). Cell culture media, trypsin, and antibiotics were purchased from HiMedia (France). Antibodies were purchased from Santa Cruz Biotechnology (Dallas, Texas, USA), and Cloud-Clone Corp. (Houston, USA). Cell Titer-Glo reagent was obtained from Promega Corp (Madison, Wisconsin, USA).

### 2.2 Cell culture

MDA-MB-231and MCF-7 cells were cultured in the L-15 medium and Dulbecco’s Modified Eagle’s medium (DMEM) respectively containing 10% FBS (Fetal bovine serum), Penicillin (100 unit/ml) and Streptomycin (100μg/ml). The cell culture was incubated at 37°C in humidified air containing 5% CO_2_.

### 2.3 Transfection

2-3^χ^10^5^ cells/well were seeded in 6 well plates one day before siRNA transfection. 25 nM of each siRNA was mixed with 100 μl Opti-MEM media. Similarly, the complex of Lipofectamine RNAi max (4 μl/each well) and Opti-MEM (100 μl) was mixed well and incubated for 5 minutes at room temperature. After that siRNA and Lipofectamine RNAiMAX complexes in Opti-MEM were mixed in 1:1 proportion and incubated for 25 minutes at room temperature. Cells were transfected with siRNA complexes and incubated at 37°C for 72 hours.

### 2.3 Cell viability assay

Cell viability was accessed by Cell Titer Glo (CTG) assay according to the manufacturer’s protocol. Briefly, MDAMB-231 cells were seeded in 96 well white culture plates at a density of 1000 cells/well in 180 μl of the medium. 20 μl of siRNA complexes of SCR, PIN1, and PTOV1 was prepared as mentioned above and reversely transfected in cells. Cells were incubated at 37°C, 5% CO_2_ for 96 hours. Following incubation, 100 μl of media was added to each well and treated with 100 μl of CTG reagent and kept on a shaker for 2 minutes. The plates were kept at room temperature for 10 minutes to stabilize the luminescence signal. Luminescence was measured using microtiter plate ELISA reader (Bio Tek, Winooski, Vermont, USA).

We also, estimated the cell viability of MCF-7 by Crystal Violet assay after transfecting cells with scramble, PIN1 and PTOV1 siRNA. Briefly, 3000 cells/well were seeded in 96 well plates and on the next day cells were transfected with the siRNAs for 72 hours. Following incubation, cells were stained with 0.4% crystal violet in 50% methanol for 30 mins. After which wells were washed with tap water to remove the excess dye. 100 μl of methanol was put in each well to dissolve the dye attached to cells on a rocker for 20 mins. Then the absorbance was recorded using microtiter plate ELISA reader.

### 2.4 Colony formation assay

MDA-MB-231 and MCF-7 cells were transfected with siRNAs for 48 hours. Then, they were collected and 1000 cells/well were seeded in 6 well plates. Cells were allowed to grow for two weeks till the colonies could be observed. The colonies were fixed with 3.7% formaldehyde for 30 minutes and stained with 0.4% crystal violet. Cells were washed with DPBS to remove excess dye. The plate was left to air dry and the colonies were counted using Image J software.

### 2.5 Wound healing assay

3^χ^10^5^ cells/well were seeded in 6 well plates and transfected with siRNAs. A scratch was made with 10 μl tip on the monolayer of cells at 72 hours post-transfection. Then they were washed with DPBS to remove detached cells. The images were captured using an inverted microscope at 0, 6, and 12 hours. The area of the wound was calculated using Image J software

### 2.6 Cell cycle analysis

2.5^χ^10^5^ cells/well were seeded in 6 well plates one day before transfection. After which they were transfected with siRNAs and incubated for 48 hours. Following the incubation, cells were harvested and fixed with ice-cold 70% ethanol and kept overnight at 4°C. Next, they were washed with DPBS, collected, and centrifuged at 1500 rpm for 5 minutes, after which they treated with RNase A (50 U/ml) for 1 hour and stained with Propidium iodide (PI) (20 μg/ml). Cells were further subjected to cell cycle analysis by flow cytometry (BD FACS Verse™).

### 2.7 Hoechst staining

1.5^χ^10^5^ cells were seeded in 6 well plates. Next day, cells were transfected with siRNAs and incubated for 72 hours. After which they were stained with Hoechst 33342 stain (2μg) and kept in the dark for 15 minutes. Next, they were washed with DPBS and the images were captured using Nikon fluorescent microscope.

### 2.8 Measurement of the intracellular ROS production

Intracellular ROS generation was detected using fluoroprobe CM-H2DCFDA (Invitrogen). 1.5^χ^10^5^ cells were seeded in 6 well plates and transfected with siRNAs for 72 hours. Cells were treated with 10 μM CM-H2DCFDA and kept in the dark at 37^0^C for 30 minutes. Then, they were stained with Hoechst 33342 and incubated for 15 minutes in the dark at room temperature. The wells were washed with DPBS and the images were captured using a fluorescence microscope.

### 2.9 RNA extraction, cDNA preparation, and real time PCR

2^χ^10^5^ cells were seeded in 6 well plates, transfected with siRNAs and incubated for 72 hours. Then cells were harvested for RNA isolation using TRIzol lysis reagent following the manufacturer’s protocol. 2μg of RNA was used for cDNA synthesis using RevertAid First strand cDNA synthesis kit (Thermo Fisher Scientific). Real time PCR was carried out using iTaq SYBR green mix (Biorad) in ABI 700, (Invitrogen). PCR cycle condition was as shown below.

Hold stage 95°c (10min), PCR Stage (40 cycles) 95°c (15 sec),60°c (1min) and melt curve stage 95°c (15 sec),60°c (1min),95°c (15 sec). Sequences of primers for Real Time PCR are; *PIN1* forward primer sequence 5’-TTTGAAGACGCCTCGTTTGC-3’, reverse primer sequence 5’-GTGCGGAGGATGATGTGGAT-3’, *PTOV1* forward primer sequence 5’-AGACACTGAAGAGCCTGTGC-3’, reverse primer sequence 5’-CGTTGACAAAGTTGCCCTGG-3’, *p21* forward primer sequence 5’CTGCCCAAGCTCTACCTTCC-3’, reverse primer sequence 5’-CGAGGCACAAGGGTACAAGA-3’, *p27* forward primer sequence 5’-TGCAACCGACGATTCTTCTACTCAA-3’, reverse primer sequence 5’-CAAGCAGTGATGTATCTGATAAACAAGGA-3’, *SURVIVIN* forward primer sequence 5’-GGACCACCGCATCTCTACAT-3’, reverse primer sequence 5’-GAAACACTGGGCCAAGTCTG-3’, *NANOG* forward primer sequence-5’-GGACCACCGCATCTCTACAT-3’, reverse primer sequence-5’-TGCAGAAGTGGGTTGTTTGC-3’, *CYCLINB1* forward primer sequence-5’-AGGCGAAGATCAACATGGCA-3’, reverse primer sequence-5’-AGGCGAAGATCAACATGGCA-3’, *CYCLINE1* forward primer sequence-5’-ATACTTGCTGCTTCGGCCTT-3’, reverse primer sequence-5’-TCAGTTTTGAGCTCCCCGTC-3’, *CYCLINA1* forward primer sequence-5’-GAAAATGCCTTCCCTCCAGC-3’, reverse primer sequence-5’-TGTGCCGGTGTCTACTTCAT-3’, *CDK1* forward primer sequence-5’-TACAGGTCAAGTGGTAGCCA-3’, reverse primer sequence-5’-AGCACATCCTGAAGACTGACT-3’, *mTOR* forward primer sequence-5’-GCAGAAGGTGGAGGTGTTTG-3’, reverse primer sequence-5’-CATTGACATGACCGCTAAAGAACG-3’, *SOX2* forward primer sequence-5’-TTTGTCGGAGACGGAGAAGC-3’, reverse primer sequence-5’-TAACTGTCCATGCGCTGGTT-3’, *SLUG* forward primer sequence-5’-ACGCCTCCAAAAAGCCAAAC-3’, reverse primer sequence-5’-ACTCACTCGCCCCAAAGATG-3’, *c-Myc* forward primer sequence-5’-CCCTCCACTCGGAAGGACTA-3’, reverse primer sequence-5’-GCTGGTGCATTTTCGGTTGT-3’, *RACK1* forward primer sequence-5’-ATGGGATCTCAACGAAGGCA-3’, reverse primer sequence-5’-CACACAGCCAGTAGCGGTTA-3’, *CIAP* forward primer sequence-5’-AAGGAGTCTTGCTCGTGCTG-3’, reverse primer sequence-5’AGCATCAGGCCACAACAGAA-3’, *E2F1* forward primer sequence-5’-ACTCAGCCTGGAGCAAGAAC-3’, reverse primer sequence-5’GGTGGGGAAAGGCTGATGAA-3’, *PCNA* forward primer sequence-5’-TCTGAGGGCTTCGACACCTA-3’, reverse primer sequence-5’-CATTGCCGGCGCATTTTAGT-3’, and *GAPDH* forward primer sequence-5’-GTGAACCATGAGAAGTATGACAAC-3’, reverse primer sequence-5’-CATGAGTCCTTCCACGATACC-3.

### 2.10 Western blotting

2^χ^10^5^ cells were cultured in 6 well plates. Next day, cells were transfected with siRNAs and incubated for 72 hours. Following incubation, cells were washed twice with ice-cold DPBS, and the cell lysate was prepared using sodium dodecyl sulphate (SDS) lysis buffer. The samples were heated for 5 minutes at 100°C in a dry bath, run on SDS-PAGE, and then transferred to PVDF membrane (100 volts, 1 hour). The membranes were incubated with 5% skim milk in Tris buffer saline with 0.1% Tween20 (TBST) for 2 hours at room temperature. The blots were further incubated with primary antibodies and kept overnight at 4°C on a rocker. Then the blots were washed 5 times with TBST and probed with horseradish peroxidase (HRP) conjugated secondary antibodies. The blots were exposed to X-ray film in the dark using enhanced chemiluminescence (ECL, Biorad). β actin was used as a loading control.

### 2.11 Co-Immunoprecipitation

The MDA-MB-231 cells were washed with cold PBS 2 times and and collected in PBS using a scrapper. Cells were centrifuged at 1000 rpm for 5 minutes. Cells were lysed in lysis buffer (50 mM Tris (pH 7.5), 5mM EDTA, 150 mM NaCl, 1 mM dithiothreitol, 0.01% Nonidet P-40, 0.2 mM phenylmethylsulfonyl fluoride, and 1X protease inhibitor mixture). Cells were sonicated for 10 seconds. Cells were again centrifuged for 15,000 rpm for 20 minutes at 4°C. The supernatant was collected in an Eppendorf tube. 50μl of lysate was collected for input. Remaining cell lysate was divided into two equal parts in 2 Eppendorf tube and made 1 ml using lysis buffer and labelled as Pin1 IP and normal rabbit IgG. The lysates were incubated with 2 μg of PIN1 and normal rabbit IgG in respective tubes at 4°C with gentle rolling for 12-24 hours. On the next day, homogenous solution of magnetic beads was taken in two Eppendorf tubes and placed in a magnet. The supernatant was thrown. The lysates containing antibody complex were incubated with the beads for 1hour at 4°C with gentle rolling. Again, the tubes were placed in the magnet and the supernatant was thrown. The beads were washed with 200μl of washing buffer for 5 times by gentle pipetting with ice incubation for 30 seconds in each wash. The tubes were placed in a magnet and the supernatant was thrown. This was repeated for 5 times. The samples along with input were heated with 2XSDS sample buffer for 5 minutes at 100°C. The samples were again placed in the magnet and the supernatant was collected in new tubes. The input (2% of lysate volume), PIN1 IP and normal rabbit IgG Samples were loaded on 10% SDS PAGE and subjected to electrophoresis^8^. Immunoblotting was done with PIN1 and PTOV1 antibody and developed using ECL.

### 2.12 Statistical Analysis

Results are represented as a mean ± standard deviation. The level of significance between scramble (control) groups and the tests (PIN1 and PTOV1 siRNA treated) groups have been calculated by *ONE WAY ANOVA* and students’s t-test. P < 0.05 was considered statistically significant. *, **, *** represents P< 0.05, 0.01 and 0.001 respectively.

## 3. Results

### 3.1. Silencing of PTOV1 and PIN1 suppresses the cell viability of MDA-MB-231 cells

Cells were knockdown for PIN1 and PTOV1 genes by transfecting with siRNAs and incubated for 96 hours. Cell viability was estimated by using CTG reagent which measures the level of ATP present in the sample which indicates cell viability. The cell viability of MDA-MB-231 was significantly inhibited after silencing of PIN1 and PTOV1 genes by siRNAs in MDA-MB-231 cells (Figure 1A). The result suggested that both the genes are essential for the proliferation of MDA-MB-231 cells. Similarly, we have performed crystal violet assay to test cell viability in MCF-7 cells after transfecting with siRNAs. The results showed similar effect in MCF-7 cells as well (Figure S1 A). As shown in figure 1F, western blot images of PIN1 and PTOV1 knockdown by their respective siRNA are shown.

**Figure 1.**
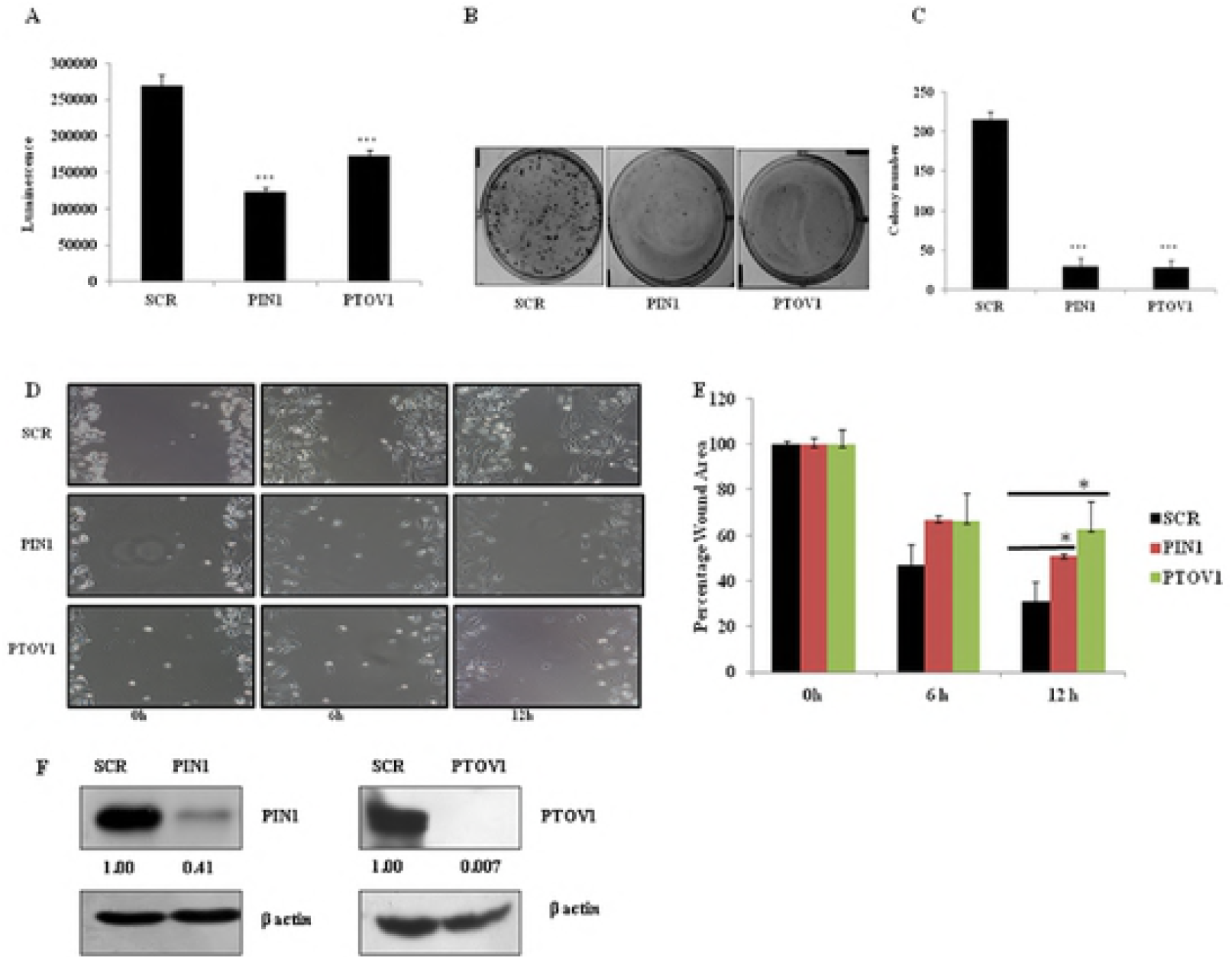
Silencing of PIN1 and PTOV1 decreases the proliferation, colony formation, and migration of MDA-MB-231 cells. (A) Cell viability was estimated by using CTG assay and luminescence signal was recorded in microtiter plate ELISA reader. (B) Representative images of the colony of cells transfected with siRNAs in 6 well plates after 2 weeks of incubation. (C) Densitometry representation of colony number using Image J software. (D) Representative images of the wound area of each treated group after 0, 6, and 12 hours. (E) Densitometry representation of Wound area using Image J software. (F) Representative image of PIN1 and PTOV1protein knockdown after transfecting cells with respective siRNAs. β- actin was used as a loading control. Data are represented as mean ± SD of three independent experiments. *** Significant difference from scramble groups (p < 0.001).

### 3.2 Silencing of PTOV1 and PIN1 inhibits colony formation of MDA-MB-231 cells

To study the long-term effect of the silencing of PIN1 and PTOV1 on cell proliferative potential, we performed colonogenic assay which is associated with tumor formation *in vivo* (11). Briefly, cells were transfected with siRNAs for 48 hours, collected, and 1000 cells per well were seeded in 6 well plates. Cells were allowed to grow for 14 days. The number and size of colonies were significantly inhibited in PIN1 and PTOV1 knockdown sample as compared to scramble (p< 0.001) (Figure 1B). Individual colonies were counted using Image J software and densitometry analysis was done (Figure 1C). Thus, silencing of both genes inhibited the proliferation and colony formation of MDA-MB-231 cells. As shown in Figure S1. B &C, similar effect has been seen in MCF-7 cells with PIN1 and PTOV1 knock down by siRNA transfection. colony numbers and size in PIN1 and PTOV1 knockdown cells were significantly inhibited as compared to scramble.

### 3.3 Silencing of PTOV1 and PIN1 attenuates the migratory potential of MDA-MB-231 cells

The migratory behavior of a cell is an important parameter in studying the invasive and metastatic potential following the treatment. Wound healing assay is a conventional technique to study the migratory behavior of a cell (12). In our experiment, we made a scratch on the monolayer of cells previously transfected with the above-mentioned siRNAs. The images were captured at 0, 6, and 12 hours using an inverted microscope. We observed that cell migration was significantly attenuated in PIN1 and PTOV1 knockdown samples as compared to scramble (Figure 1D). Quantitative analysis of the wound area using Image J software showed significant difference between scramble and PIN1 and PTOV1 knockdown cells at 12hr (p< 0.05) (Figure 1E). These results suggested that both the genes might be crucial for the migration of cells. In similar manner, we studied migration potential of MCF-7 cells and found that migration was significantly inhibited in 24 and 48 hours after knocking down of PIN1 and PTOV1 as shown in Figure S1. D and E. Thus, the result showed both PIN1 and PTOV1 genes are important for migration of MCF-7 cells.

### 3.4 Silencing of PTOV1 and PIN1 induces G2/M phase arrest of MDA-MB-231 cells

After 48 hours of knocking down of cells with PIN1 and PTOV1 siRNAs, cells were subjected to cell cycle analysis by flow cytometry. Briefly, cells were collected, fixed with ice-cold 70% ethanol and incubated overnight at 4°C. This was followed by RNase treatment and staining with Propidium Iodide (PI). Percentage distribution of the cell in different phases of the cell cycle was estimated using flow cytometry which showed that there was a significant proportion of cells in G2/M phase for PTOV1 siRNA treated groups (19.73%) as compared to that in scramble siRNA treated groups (13.266%) (p < 0.05) (Figure 2 A). The results showed that silencing of PTOV1 has an impact on cell cycle, but the effect was moderate. Knockdown of PTOV1 might block the entry of cells into the mitotic phase. We have also performed cell cycle analysis after 48 hours of knocking down of PIN1 in MDA-MB-231 cells. Result showed that PIN1 knockdown arrests cells in G2/M phase significantly showing similar effect as PTOV1[scramble (10.89%) & PIN1(17.65%)]. (Figure 2B)

**Figure 2.**
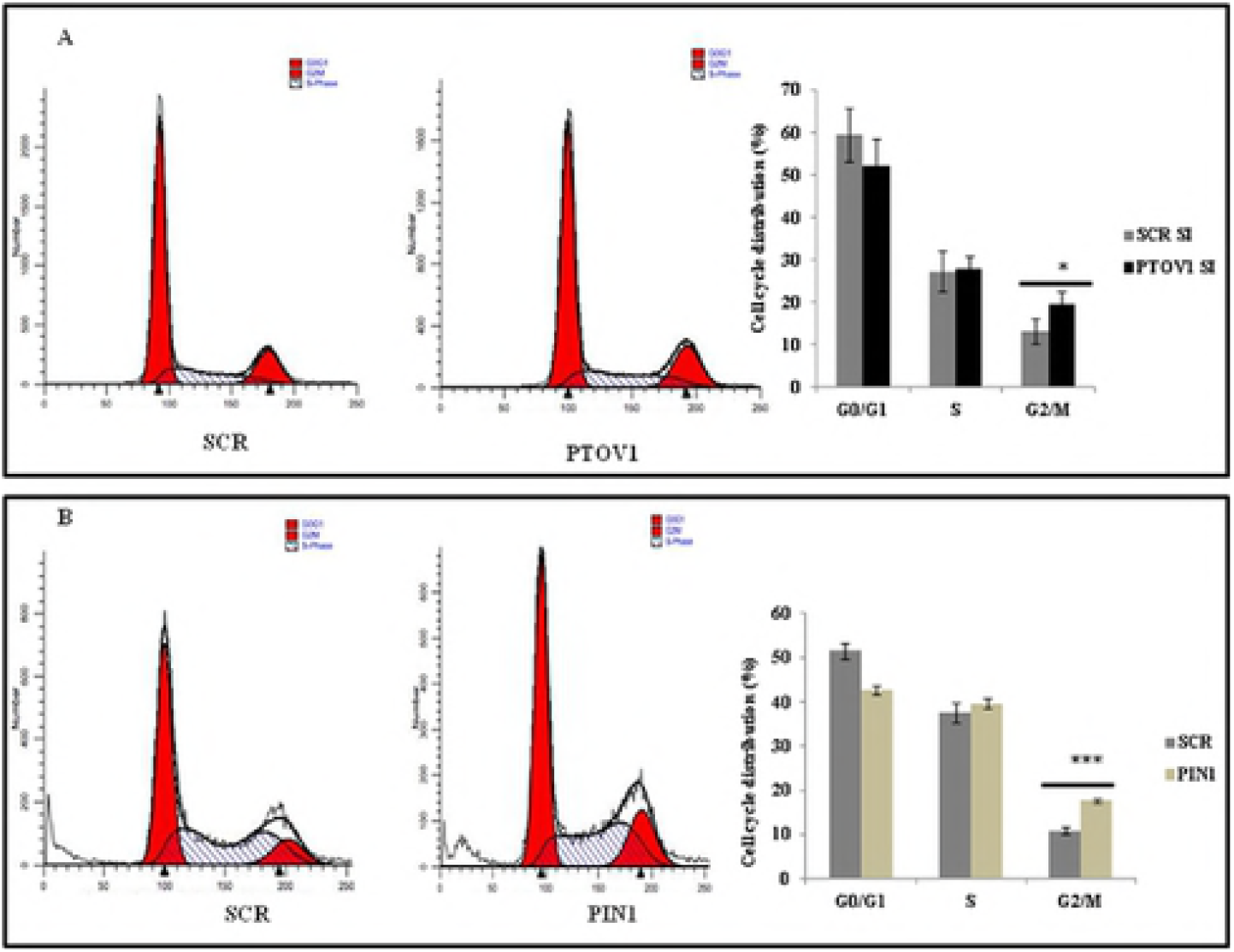
Silencing of PTOV1 and PIN1 leads to the G2/M arrest in MDA-MB-231 cells. (A) Histogram representing the cell cycle distribution of MDA-MB-231 cells transfected with PTOV1 siRNA for 48 hours (left panel) and statistical representation of percentage distribution of the population in each phase (right panel). (B) Histogram representing the cell cycle distribution of MDA-MB-231 cells transfected with PIN1siRNA for 48 hours (left panel) and statistical representation of percentage distribution of the population in each phase (right panel). Data are represented as mean ± SD of three independent experiments. *, *** Significant difference from scramble groups (p < 0.05 & 0.001).

### 3.5 Silencing of PTOV1 and PIN1 induces nuclear condensation and initiation of apoptosis

Nuclear condensation is the phenotype which indicates the initiation of apoptosis (13, 14). Cells were transfected with the siRNAs for 72 hours and then stained with Hoechst 33342 dye. The phase contrast images showed that the morphology of cells in PIN1 and PTOV1 samples were altered and not healthy as compared to untransfected and scramble and the DAPI images showed that the nuclear size was reduced in knockdown samples compared to scramble and untransfected cells. These results showed that silencing of both the genes might initiate cells towards apoptosis (Figure 3A).

### 3.6 Silencing of PTOV1and PIN1 elevates the intracellular ROS production

The level of ROS production has a determinant role on the behavior of cells. In the context of cancer, it possesses both an oncogenic property sustaining the proliferation of cancer cells as well as a tumor suppressor property when there is an aberrant increase in ROS due to any kind of stimuli (15). Thus, the search for the genes modulating the redox status of a cancer cell can be a good anti-cancer strategy (16). As shown in Figure 3B, there was an increased production of ROS in PIN1 and PTOV1 silenced cells compared to scramble siRNA and non-transfected cells. Bar graph in right panel showed the quantification of ROS production by counting number of cells producing ROS out of total number of cells. Thus, our results revealed that downregulation of PIN1 and PTOV1 modulates the redox status of MDA-MB-231 cells enhancing their death significantly.

### 3.7 Silencing of PTOV1 and PIN1 initiates the cell death of MDA-MB-231 cells by inducing apoptosis

To study the cell death mechanism initiated by knockdown of PIN1 and PTOV1, western blot analysis was carried out to check the expression of apoptosis-related genes. There was inhibition of anti-apoptotic Bcl-2 (Figure 4A), Bcl-xL (Figure 4B), and up-regulation of pro-apoptotic BAX (Figure 4C). These results clearly suggested that inhibition of both the genes initiate cell death through apoptosis. Also, we found the enhanced expression of autophagy markers LC3 (Figure 4D) and Beclin-1(Figure 4E) after PTOV1 (Beclin-1 was unaffected by PIN1 knockdown figure 4F), These results demonstrated that PTOV1 silencing also induced cell death by autophagy.

### 3.8 Silencing of PTOV1 and PIN1 shares regulatory networks and affect diverse cellular functions at the transcriptional level

mRNA expression of the genes affecting cell proliferation, cell cycle, apoptosis, and metastasis was studied after knockdown of cells with siRNAs for 72 hours. As shown in Figure 5A & B, the bar graph showed the relative expression of mRNAs in PIN1 and PTOV1 siRNA transfected samples compared to scramble samples respectively. The value for the expression of genes in scramble sample is considered to be 1.00.

Study of the regulation of various genes by PIN1 and PTOV1 at transcriptional level mentioned in Table No. 1, we realized that knockdown of both the genes shared similar expression profile and affect the major functions required for cancer cell progression. The expression was downregulated for cell proliferation, cell cycle, and metastasis-related genes while cell cycle inhibitor p27 and p21 were upregulated in PIN1 and PTOV1 siRNA transfected samples respectively. Our results suggest that a wide range of genes which are involved in cell proliferation, cell cycle, metastasis, and apoptosis are regulated by PIN1 and PTOV1 at the transcriptional level significantly.

**Table No. 1.**
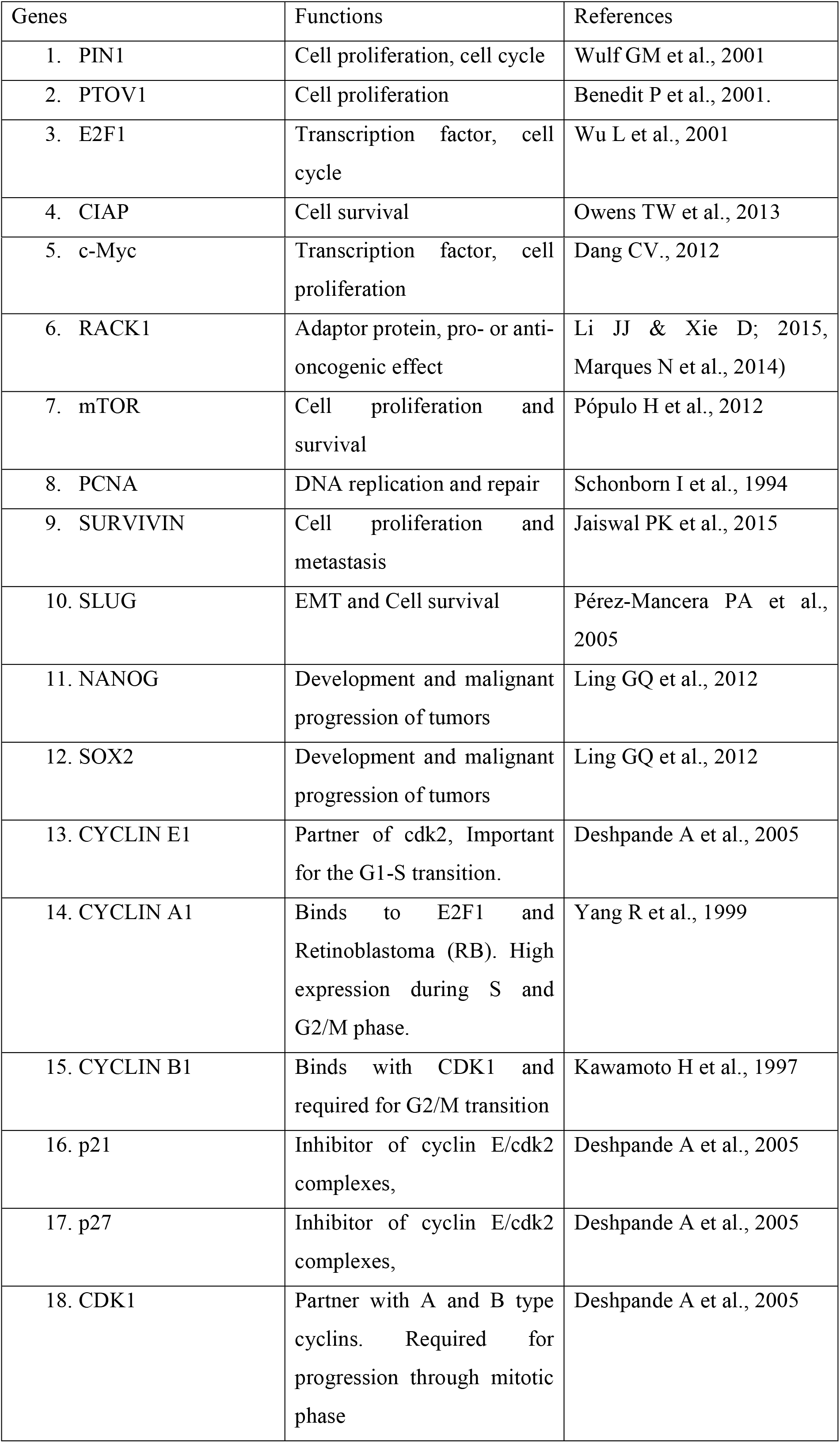
The genes used for the mRNA expression analysis with their cellular functions are listed in the table below.

### 3.9 PTOV1 interacts with PIN1 and share the regulatory network of multiple oncogenic signaling pathways at the translational level

We have already reported that PIN1 interacts with PTOV1 in PC-3 cells (8). We have performed Co-IP with PIN1 IP to see the interaction of PIN1 and PTOV1 in MDA-MB-231 cells. As shown in figure 6A. (i) PTOV1 is enrich in PIN1 complex. Furthermore, we analyzed the common regulatory network of both the genes at the protein level, western blots were performed after transfecting cells with siRNAs for 72 hours. Knockdown of PIN1 inhibited the expression of PTOV1 but there was no change in the expression of PIN1 protein level with PTOV1 knockdown (Figure 6A. ii & iii). Cells were also treated with Juglone (5μM), a PIN1 inhibitor, for 48 hours. The expression of both PIN1 and PTOV1 was inhibited significantly (Figure 6A. iv). These results suggested that PTOV1 contributes for PIN1 mediated MDA-MB-231 cell proliferation. Moreover, the overexpression of PIN1 with PIN1-XPRESS increased the expression of PTOV1 more than two folds (Figure 6A. v). These results supported our previous finding that PTOV1 and PIN1 interaction play an important role in cancer cell proliferation (8).

PIN1 and PTOV1 silencing inhibited the expression of β-catenin (Figure 6B. i) which is involved regulates the expression of both Cyclin D1 and c-Myc (Figure 6B. ii, & iii). We have earlier shown that PTOV1 regulates the PIN1 substrate c-Jun phosphorylation (8). These data supported our finding that PIN1 and PTOV1 contribute to oncogenesis of breast cancer cells targeting Cyclin D1 and c-Myc expression. PTOV1 interacts with the RACK1 to bind with the 40s ribosomes during translation initiation (10). RACK1 has earlier been shown to be the regulator of cell signaling, migration and ribosomal functioning (29–31). In our experiment, we found that PIN1 and PTOV1 knockdown suppressed the expression of RACK1. Thus, our results showed that PIN1 is required for PTOV1 mediated translational machinery through RACK1 (Figure 6C. i). We also checked the expression of Cyclin B1, a G2/M transition regulatory protein and found that its expression was inhibited by siRNAs (Figure 6C. ii). This result supported the role of PTOV1 in cell cycle regulation. The expression of MAPK8 (c-Jun N-terminal kinase) which phosphorylates c-Jun (Figure 6C. iii) was also inhibited. ERK is already known to be the substrate of PIN1 (32). Decreased phosphorylated ERK by PTOV1 downregulation suggested that PIN1 and PTOV1 share the similar expression profile of MAPK pathway (Figure 6C. iv). These results showed the mechanisms of the PIN1 and PTOV1 cascade involvement in breast cancer cell proliferation via β-catenin and MAPK pathway.

## 4. Discussion

PTOV1 expression predicts the poor prognosis in multiple tumors (9, 10, 33, 34). Studies on the interacting partners and the proteins that share the close regulatory network with PTOV1 has helped to elucidate the cellular functions. Our study has taken the first step in establishing the functional relationship between PIN1 and PTOV1. On the assumption that functionally related genes show a similar expression profile and could fall in the same pathway, in our study we have used the RNAi mediated approach to knock down the genes and explore the different parameters of cell transformation such as cell proliferation, migration, cell cycle, autophagy and apoptosis that are required to draw the correlation between the genes. Our earlier finding showed that PTOV1 physically interacts with PIN1. PTOV1 also regulates the expression of PIN1 substrate c-Jun phosphorylation which is a major transcription factor involved in cell proliferation, migration, and apoptosis (8).

Cell viability and colony formation of MDA-MB-231 and MCF-7 cells were significantly reduced after PIN1 and PTOV1 knockdown, which suggests that both the genes have a strong impact on the growth of cells. Silencing of PIN1 and PTOV1 strongly inhibited the migration of cells. The role of PTOV1 in cell cycle progression has been studied in prostate cancer cells(33). There are numerous reports on the role of PIN1 in cell cycle (34, 35). Here, we have shown that PIN1 and PTOV1 knockdown arrests breast cancer cell MDA-MB-231 in the G2/M phase of the cell cycle before entry into the mitosis where rapid synthesis of proteins is required for cell division. This supports the role of PTOV1 in protein translation.

Modulation of redox status can be employed for the killing of cancer cells (16). We found that PIN1 and PTOV1 both hold promising target to change the redox status of cancer cells. The balance between cell proliferation and regression of tumor determines the efficacy of radiotherapy, chemotherapy or hormonal therapy, which is governed, in part, by the repertoire of pro-apoptotic and anti-apoptotic molecules (36-40). Here, we have studied the expression of Bcl-2 and Bcl-xL (anti-apoptotic) and BAX (pro-apoptotic) after knocking down of our target genes. Downregulation of Bcl-2, Bcl-xL and up-regulation of BAX confirmed that cells were driven towards apoptosis after the knocking down of PIN1 and PTOV1 respectively. Therapeutic targeting for autophagy induction is a newer form of cell death in cancer which depends on cell type (41). Knockdown of PTOV1 increased the expression of autophagy markers Beclin-1 and LC3, suggesting that cell death also induced by autophagy.

Both PIN1 and PTOV1 knockdown regulates multiple genes involved in MDA-MB-231 cell proliferation, cell cycle, metastasis, and apoptosis in their transcriptional level. Western blot analysis was performed to study the expression of different proteins related to cell transformation. First, we studied the interaction between PIN1 and PTOV1in MDA-MB-231 cells by co-immunoprecipitation. We found that PTOV1 was enriched in the PIN1 complex. Silencing of PIN1 downregulated the expression of PTOV1 but not vice-versa. Treatment with Juglone completely blocked the expression of PTOV1. Also, the ectopic expression of PIN1 increased the expression of PTOV1. Taken together, all these results show that PIN1 is upstream of PTOV1 in the signaling cascade. Activation of Wnt/β-catenin is a good predictor with poor prognosis in breast cancer (42, 43). PIN1 and PTOV1 downregulation inhibited the expression of β-catenin. This result showed that PIN1 and PTOV1 might be linked for the activation of Wnt/β-catenin signaling pathway. Our results show that PIN1 and PTOV1 target Cyclin D1 and c-Myc. Extracellular signal-regulated kinases (ERKs), one of the members of mitogen-activated protein kinases (MAPKs), are the substrates for proline-directed phosphorylation by PIN1(32). Silencing of PTOV1 also abolished the phosphorylation of ERK. It can be speculated that PTOV1 may be required for the PIN1 mediated phosphorylation of ERK. RACK1 has earlier been shown to interact with PTOV1 which regulates translation of c-Jun in PC-3 cells (10). Our previous finding showed that PTOV1 is a novel regulator of PIN1 mediated c-Jun phosphorylation and PTOV1 interacts with PIN1 (8). Here, we have shown that RACK1 is regulated by both PIN1 and PTOV1. Summing up all these results, it can be deduced that PTOV1 is contributing for PIN1 mediated cell transformation.

In conclusion, we have demonstrated that PIN1 and PTOV1 show similar expression profiles of a cohort of genes by RNAi silencing in MDA-MB-231 cells. Functional validation of their relationship was found to be both at transcriptional and translational levels and showed their effect on cell viability, colony forming potential, cell migration, cell cycle, and metastasis.PIN1 and PTOV1 drives the expression of Cyclin D1 c-Myc & β-catenin. Also silencing of genes affect the proteins related to MAPK pathway. Both of genes affect the overall survival of MDA-MB-231 cells by targeting BAX, Bcl-2, and Bcl-xL as shown in figure 7. Thus, both PIN1 and PTOV1 drives the common genes regulation to impart oncogenic phenotype of MDA-MB-231 cells, thus PIN1-PTOV1 complex could be a potential target for future cancer therapy.

## Acknowledgement

This project is funded by The South Asian University Start Up Fund

## Conflict of Interest

Authors declare no conflicts of interest for this article

---

**Figure 3. Silencing of PIN1 and PTOV1 alters cell morphology, apoptosis, and intracellular ROS production.** (A) Representative images of cells transfected with the siRNAs after 72 hours and stained with Hoechst 33342. (B) Microscopic evaluation of intracellular ROS production in each group after 72 hours using CM-H2DCFDA. Images were captured using Nikon fluorescent microscope, right panel shows the graphical representation of the ROS production in different set of experiments. *** represents the significant difference from scramble (p< 0.001).

**Figure 4.** Silencing of PIN1 and PTOV1 drives cells towards apoptosis and PTOV1 knockdown further induces autophagy. Western blot analysis of (A) Bcl-2, (B) Bcl-xL, and (C) BAX after 72 hours of transfection. Expression analysis of autophagy markers (D) LC3, and (E) Beclin-1after transfecting cells with scramble and PTOV1 siRNA for 72 hours (F) Beclin-1 after transfecting cells with respective scramble and PIN1 siRNAs. The density was calculated by image J software. β- actin was used as a loading control. The experiments were performed in triplicates.

**Figure 5. Silencing of PIN1 and PTOV1 shares similar expression profile for the majority of genes at the transcriptional level.** (A) The upper panel represents the expression of mRNA after transfecting cells with PIN1 siRNA for 72 hours. (B) The lower panel represents the expression of mRNA after knockdown with PTOV1 siRNA for 72 hours. Relative expression was calculated using delta-delta CT method. The expression in scramble group was considered 1.00. Experiments were performed in triplicates. Data are represented as mean ± SD.*, **, *** represents significant difference from the scramble with p< 0.05, 0.01 and 0.001 respectively.

**Figure 6. Silencing of PIN1 and PTOV1 cooperatively targets multiple proteins and shares similar expression status at the translational level.** A. (i) PIN1 and PTOV1 interaction studied by co-immunoprecipitation in MDA-MB-231 cells. (ii& iii) Western blot analysis of PIN1 and PTOV1 after transfecting cells with siRNAs for 72 hours, (iv) with Juglone (5 μM) for 48 hours, and (v) Overexpression with PIN1-XPRESS vector. Expression analysis of B. (i) β- catenin, (ii) Cyclin D1, (iii) c-Myc, & further analysis of expression of C. (i) RACK1, (ii) Cyclin B1, (iii) MAPK8, and (iv) p-ERK after 72 hours of siRNA transfection. The densitometry was calculated by using image J software. β-actin was used as a loading control. The experiments were performed in triplicates.

## References

1. Ghoncheh M, Pournamdar Z, Salehiniya H, et al. Incidence and Mortality and Epidemiology of Breast Cancer in the World. Asian Pac J Cancer Prev.17: 43–6, 2016.

2. Clegg LX, Reichman ME, Miller BA, Hankey BF, Singh GK, Lin YD, Goodman MT, Lynch CF, Schwartz SM, Chen VW, Bernstein L, et al. Impact of socioeconomic status on cancer incidence and stage at diagnosis: selected findings from the surveillance, epidemiology, and end results: National Longitudinal Mortality Study. Cancer causes & control. 20: 417–435, 2009.

3. Berry DA, Cronin KA, Plevritis SK, Fryback DG, Clarke L, Zelen M, Mandelblatt JS, Yakovlev AY, Habbema JDF, Feuer EJ, et al. Effect of screening and adjuvant therapy on mortality from breast cancer. New England Journal of Medicine. 353: 1784–1792, 2005.

4. Ravdin PM, Cronin KA, Howlader N, Berg CD, Chlebowski RT, Feuer EJ, Edwards BK, Berry DA et al. The decrease in breast-cancer incidence in 2003 in the United States. New England Journal of Medicine.356 :1670–4, 2007.

5. Garner AL, Janda KD et al. Protein–protein interactions and cancer: targeting the central dogma. Current topics in medicinal chemistry.11 :258–80, 2011.

6. Ivanov AA, Khuri FR, Fu H et al. Targeting protein-protein interactions as an anticancer strategy. Trends in pharmacological sciences. 34 :393–400, 2013.

7. Wulf GM, Ryo A, Wulf GG, Lee SW, Niu T, Petkova V, Lu KP et al. Pin1 is overexpressed in breast cancer and cooperates with Ras signaling in increasing the transcriptional activity of c-Jun towards cyclin D1. The EMBO journal. 20 :3459–72, 2001.

8. Pokharel YR, Saarela J, Szwajda A, Rupp C, Rokka A, Karna SK, Teittinen K, Corthals G, Kallioniemi O, Wennerberg K, Aittokallio T et al. Relevance rank platform (RRP) for functional filtering of high content protein-protein interaction data. Molecular & Cellular Proteomics. 14(12):3274–83, 2015.

9. Benedit P, Paciucci R, Thomson TM, Valeri M, Nadal M, Càceres C, de Torres I, Estivill X, Lozano JJ, Morote J, Reventós J et al. PTOV1, a novel protein overexpressed in prostate cancer containing a new class of protein homology blocks. Oncogene. 2:1455, 2001.

10. Marques N, Sesé M, Cánovas V, Valente F, Bermudo R, De Torres I, Fernandez Y, Abasolo I, Fernandez PL, Contreras H, Castellon E et al. Regulation of protein translation and c-Jun expression by prostate tumor overexpressed 1. Oncogene.; 33:1124, 2014.

11. Tiang JM, Butcher NJ, Minchin RF et al. Small molecule inhibition of arylamine N-acetyltransferase Type I inhibits proliferation and invasiveness of MDA-MB-231 breast cancer cells. Biochemical and biophysical research communications. 39: 95–100, 2010.

12. Rodriguez LG, Wu X, Guan JL et al. Wound-healing assay. Methods Mol Biol. 294: 23–9, 2005.

13. Saraste A, Pulkki K et al. Morphologic and biochemical hallmarks of apoptosis. Cardiovascular research. 45:528–37, 2000

14. Crowley LC, Marfell BJ, Waterhouse NJ et al. Analyzing cell death by nuclear staining with Hoechst 33342. Cold Spring Harbor Protocols. 2016:pdb-rot087205, 2016.

15. Wang J, Yi J et al. Cancer cell killing via ROS: to increase or decrease, that is the question. Cancer biology & therapy. 7:1875–84, 2008

16. Gurer-Orhan H, Ince E, Suzen S, Saso L. The Role of Oxidative Stress Modulators In Breast Cancer. inflammation. 25(33):4084–4101, 2018.

17. Wu L, Timmers C, Maiti B, Saavedra HI, Sang L, Chong GT, Nuckolls F, Giangrande P, Wright FA, Field SJ, Greenberg ME et al. The E2F1–3 transcription factors are essential for cellular proliferation. Nature. (6862):457, 2001.

18. Owens TW, Gilmore AP, Streuli CH, Foster FM. Inhibitor of apoptosis proteins: promising targets for cancer therapy. Journal of carcinogenesis & mutagenesis. Suppl 14.pii: S14-004, 2013.

19. Dang CV. MYC on the path to cancer. Cell.149: 22–35, 2012.

20. Li JJ, Xie D et al. RACK1, a versatile hub in cancer. Oncogene. 34:1890. 2015

21. Pópulo H, Lopes JM, Soares P et al. The mTOR signalling pathway in human cancer. International journal of molecular sciences. 13: 1886–918, 2012.

22. Schönborn I, Minguillon C, Möhner M, Ebeling K et al. PCNA as a potential prognostic marker in breast cancer. The Breast.3: 97–102, 1994.

23. Jaiswal PK, Goel A, Mittal RD et al. Survivin: A molecular biomarker in cancer. The Indian journal of medical research. 141:389, 2015.

24. Perez-Mancera PA, Gonzalez-Herrero I, Pérez-Caro M, Gutierrez-Cianca N, Flores T, Gutierrez-Adan A, Pintado B, Sánchez-Martín M, Sánchez-García I et al. SLUG in cancer development. Oncogene. 24:3073, 2005.

25. Ling GQ, Chen DB, Wang BQ, Zhang LS et al. Expression of the pluripotency markers Oct3/4, Nanog and Sox2 in human breast cancer cell lines. Oncology letters. 4: 1264–8, 2012.

26. Deshpande A, Sicinski P, Hinds PW et al. Cyclins and cdks in development and cancer: a perspective. Oncogene. 24:2909, 2005.

27. Yang R, Müller C, Huynh V, Fung YK, Yee AS, Koeffler HP et al. Functions of cyclin A1 in the cell cycle and its interactions with transcription factor E2F-1 and the Rb family of proteins. Molecular and cellular biology. 19: 2400–7, 1999.

28. Kawamoto H, Koizumi H, Uchikoshi T et al. Expression of the G2-M checkpoint regulators cyclin B1 and cdc2 in nonmalignant and malignant human breast lesions: immunocytochemical and quantitative image analyses. The American journal of pathology.150:15, 1997.

29. Sengupta J, Nilsson J, Gursky R, Spahn CM, Nissen P, Frank J. Identification of the versatile scaffold protein RACK1 on the eukaryotic ribosome by cryo-EM. Nature Structural and Molecular Biology.11:957, 2004.

30. Buensuceso CS, Woodside D, Huff JL, Plopper GE, O’Toole TE et al. The WD protein Rack1 mediates protein kinase C and integrin-dependent cell migration. Journal of cell science.114: 1691–8, 2001.

31. Coyle SM, Gilbert WV, Doudna JA et al. Direct link between RACK1 function and localization at the ribosome in vivo. Molecular and cellular biology. 29: 1626–34, 2009.

32. Lu Z, Hunter T et al. Prolyl isomerase Pin1 in cancer. Cell research. 24:1033, 2014.

33. Santamaría A, Fernández PL, Farré X, Benedit P, Reventós J, Morote J, Paciucci R, Thomson TM et al. PTOV-1, a novel protein overexpressed in prostate cancer, shuttles between the cytoplasm and the nucleus and promotes entry into the S phase of the cell division cycle. The American journal of pathology. 162: 897–905, 2003.

34. Lin CH, Li HY, Lee YC, Calkins MJ, Lee KH, Yang CN, Lu PJ et al. Landscape of Pin1 in the cell cycle. Experimental Biology and Medicine. 240: 403–8, 2015.

35. Yeh ES, Means AR et al. PIN1, the cell cycle and cancer. Nature Reviews Cancer.7(5):381, 2007.

36. Reed JC. Dysregulation of apoptosis in cancer. Journal of clinical oncology. 17:2941, 1999.

37. Tamm I, Schriever F, Dörken B et al. Apoptosis: implications of basic research for clinical oncology. The lancet oncology. 2: 33–42, 2001.

38. Hickman JA. Apoptosis induced by anticancer drugs. Cancer and Metastasis Reviews.11: 121–39, 1992.

39. Verheij M, Bartelink H et al. Radiation-induced apoptosis. Cell and tissue research. 30:133–42, 2000.

40. Ellis PA, Saccani-Jotti G, Clarke R, Johnston SR, Anderson E, Howell A, A’hern R, Salter J, Detre S, Nicholson R, Robertson J et al. Induction of apoptosis by tamoxifen and ICI 182780 in primary breast cancer. International Journal of Cancer. 72: 608–13, 1997.

41. Gump JM, Thorburn A et al. Autophagy and apoptosis: what is the connection? Trends in cell biology. 21: 387–92, 2011.

42. Zardawi SJ, O’Toole SA, Sutherland RL, Musgrove EA et al. Dysregulation of Hedgehog, Wnt and Notch signalling pathways in breast cancer. Histology and histopathology. 24: 385–98, 2009.

43. López-Knowles E, Zardawi SJ, McNeil CM, Millar EK, Crea P, Musgrove EA, Sutherland RL, O’Toole SA et al. Cytoplasmic localization of β-catenin is a marker of poor outcome in breast cancer patients. Cancer Epidemiology and Prevention Biomarkers.19:301–9, 2010.

